# Growth is required for perception of water availability to pattern plant root branches

**DOI:** 10.1101/097758

**Authors:** Neil E. Robbins, José R. Dinneny

## Abstract

Water availability is a potent regulator of development in plants and acts as a positional cue to induce root branching through a process termed hydropatterning. The mechanism by which roots perceive the spatial distribution of water to position lateral branches is unknown. Here we reveal that a root's developmental competence for hydropatterning is limited to the root tip, where tissue growth occurs. Mathematical modeling suggests that water uptake during growth creates spatial biases in tissue water potential, and we show that these gradients predict the position of future lateral branches. By altering growth dynamics with exogenous chemical and environmental treatments, we demonstrate that growth is necessary to allow roots to distinguish environments with relatively high or low water availability and pattern branching accordingly. Furthermore, we show that these cues regulate a number of other physiologically important pathways. Our work supports a sense-by-growth mechanism governing lateral root hydropatterning, in which water availability cues are rendered interpretable through growth-sustained water movement.

---

Water availability is a potent regulator of development in plants and acts as a positional cue to induce root branching through a process termed hydropatterning^1,2^. The mechanism by which roots perceive the spatial distribution of water to position lateral branches is unknown. Here we reveal that a root’s developmental competence for hydropatterning is limited to the root tip, where tissue growth occurs. Mathematical modeling suggests that water uptake during growth creates spatial biases in tissue water potential, and we show that these gradients predict the position of future lateral branches. By altering growth dynamics with exogenous chemical and environmental treatments, we demonstrate that growth is necessary to allow roots to distinguish environments with relatively high or low water availability and pattern branching accordingly. Furthermore, we show that these cues regulate a number of other physiologically important pathways. Our work supports a sense-by-growth mechanism governing lateral root hydropatterning, in which water availability cues are rendered interpretable through growth-sustained water movement.

Lateral root development is activated in tissues contacting a water source, such as moist soil or agar media, and inhibited in tissues exposed to environments with low water availability, such as air (Fig. 1a-c)^1,2^. Previous work had suggested that competence to respond to hydropatterning cues was limited to the root tip^1^. Roots of all vascular plants grow from their tips and new tissues form through spatially separated processes of cell division and anisotropic cell expansion, which primarily occur in the meristem and elongation zones, respectively. We reasoned that a clearer understanding of where the root is competent to respond to such environmental cues would help to elucidate the mechanisms by which the plant spatially differentiates sources of water. We utilized roots of *Zea mays* (maize) to define the hydropatterning-competent zone, as it provided experimental advantages compared to Arabidopsis due to the higher density of lateral roots relative to the size of the different developmental zones (~7-10 lateral roots/cm primary root in maize vs ~1-3 in Arabidopsis) (Extended Data Fig. 1). In addition, its larger diameter facilitated the use of micromanipulation and micro-dissection experimental approaches that could be used for further characterization of the process (1-mm diameter in maize vs 0.1 mm in Arabidopsis).

**Figure 1.**
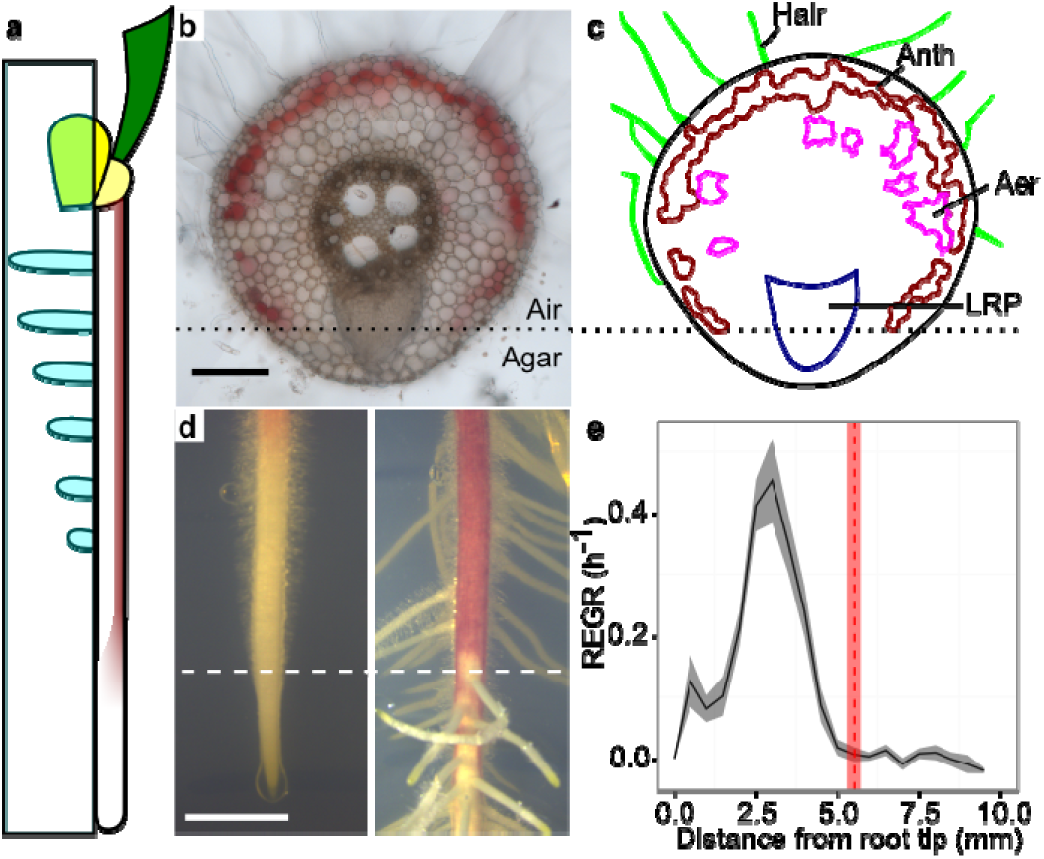
Developmental competence to respond to water availability is limited to the root tip. **a**, Diagram of maize seedling grown along agar media. Contact with agar (cyan box) locally activates lateral root development. **b-c**, Radial section of primary root (**b**) and diagram highlighting environmentally regulated anatomical features (**c**). Hair, root hair; Anth, anthocyanin; Aer, aerenchyma; LRP, lateral root primordium. Scale bar, 250 μm. **d**, Air side of primary root immediately after application of agar sheet (left) and following lateral root emergence 3 days later (right). Dashed line, boundary between competent and fixed zones. Scale bar, 5 mm. **e**, Average relative elemental growth rate (REGR) (black, n = 38 seedlings) and position of competent/fixed-zone boundary (red, n = 47 seedlings). Shaded regions, standard error of the mean. Measurements are averages of 3 experimental replicates.

We applied an agar sheet to the air-exposed side of the root and observed the pattern of lateral root development after several days, which allowed us to determine where along the length of the root the tissues maintained or lost competence to respond to this new water cue (Extended Data Fig. 1a). Strikingly, we observed that competence to respond to the agar stimulus was clearly differentiated along the length of the root, with a distinct boundary separating responsive and unresponsive regions (Fig. 1d). We refer to tissues in the rootward direction of this boundary as the competent zone, and those in the shootward direction as the fixed zone. Our results placed the boundary within published ranges of the root growth zone^3,4^; quantification of local tissue expansion rates via kinematic growth analysis^5,6^ showed that it lay just outside of this region (Fig. 1e). No significant difference was found between the measured longitudinal positions on the root where competence was lost and where growth ceased (p = 0.9, mean difference ± standard error = −0.03 ± 0.25 mm), indicating a strong correlation between these two physiological states. Indeed, past work on the early patterning of lateral root founder cells in Arabidopsis showed that this process is controlled through oscillating changes in auxin signaling that positionally correlates with the end of the growth zone^7^.

Loosening of plant cell walls allows for water uptake and growth^2,8,9^. Although the air- and agar-contacting sides of a root in our experimental system have differential access to available water, there was no obvious sign of differential growth between these regions, indicating that rates of expansion and therefore water uptake were equal on the two sides. Since all available water resides in the agar media, water must move across the root radial axis in order to sustain cell expansion in air-exposed tissues. Differences in water potential, or the chemical potential energy of water, serve as a driving force for water movement and determine its directionality: movement occurs from regions of high to low potential^2,10,11^. Thus, we predicted that a growth-sustained water potential gradient would occur in the root tip, which could provide physical cues important in hydropatterning.

Growth-sustained water potential gradients have been proposed and empirically measured in the literature^12–15^, but a role in development has not yet been established. We asked to what extent a gradient existed in the competent zone, and whether it had any impact on lateral root patterning. We created a mathematical model to estimate the extent to which growth could modulate the water potential of root tissues in our culture conditions (Extended Data Fig. 2). Briefly, water potentials were estimated based on our empirical kinematic growth measurements, which dictate the volume of water that must enter the tissue to allow for a change in volume, and values taken from the literature for hydraulic conductivity, a measure of resistance to water movement^2,10,16^.

As a proof-of-concept test for the ability of this method to accurately estimate tissue water potential, we applied the model to *Glycine max* hypocotyls, for which empirical measurements of tissue water potential are available^13^ (Extended Data Fig. 3). Accuracy of model predictions depended largely on the value used for tissue hydraulic conductivity, with highest accuracy obtained at a value within the range of those previously reported^17^. Discrepancies between overall profiles of empirical and estimated water potentials hinted at tissue-specific variation in conductivity not taken into account by the model, which assumes uniform conductivity. The absence of higher-resolution measurements of tissue hydraulic conductivity, and the experimental challenges associated with such measurements, make bulk-tissue values a necessary approximation.

We applied the model to maize roots growing along an agar medium. Local tissue water potentials were predicted to decrease as local growth rate increased, with the largest decreases predicted in tissues most distal to the external water source, thus generating a differential across the radial axis (Fig. 2a). Notably, all tissues approached water potential equilibrium after growth ceased, demonstrating the necessity of growth for generating potential gradients. These results suggest that substantial differentials in water potential exist between air- and agar-contacting tissues in the competent zone (peak differential in the epidermis = −0.60 MPa, cortex = −0.27 MPa).

**Figure 2.**
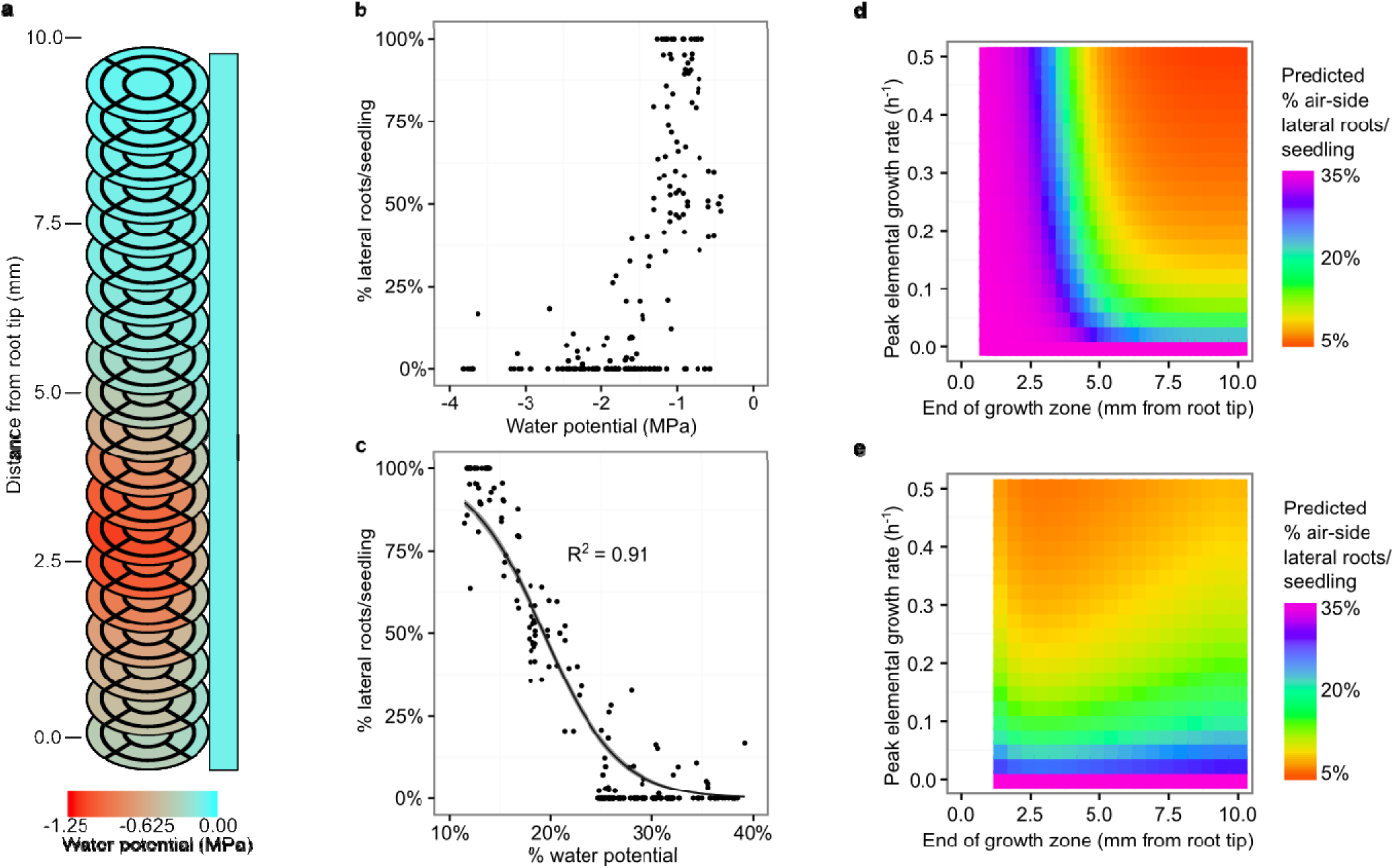
Growth-sustained water potentials are predictive of lateral root patterning in a mathematical model. **a**, Heat map of estimated tissue water potentials for maize primary root grown with a unilateral source of water (rectangle). Each disc represents one 0.5-mm root segment. Estimates were derived from empirical growth curve in Fig. 1d. **b-c**, Plots of raw water potential estimates (**b**) and normalized values (**c**) against lateral root distributions for primary roots treated with mannitol (Extended Data Fig. 4). Curve and shaded region, mean ± standard error of best-fit line for zero-one inflated beta regression model. R^2^, pseudo-R^2^ value. **d-e**, Predicted frequency of air-side lateral root initiation with varied growth zone parameter values. Competent zone was set as a 1.5-mm window that ended 5.5 mm from the root tip (**d**) or at the end of the growth zone in each simulation (**e**). Start of growth zone was fixed at 0 mm from the root tip in all simulations. Predictions were generated for primary roots grown between two agar sheets.

In order to determine whether such potentials play a role in hydropatterning, we first tested whether a relationship existed between tissue water potential and lateral root development. Kinematic analysis of root growth rate was quantified in seedlings grown between two agar sheets containing differing concentrations of mannitol, which allowed external water availability to be varied over a wide range of values (Extended Data Fig. 4a-b). These data were used to estimate competent-zone tissue water potential based on our model (Extended Data Fig. 5). Lateral root patterning was measured in these same plants across the different air- and agar-contacting sides of the root (Extended Data Fig. 4c). Interestingly, while absolute water potential did not show an obvious relationship with lateral root patterning, relative water potential (normalized between domains of the same root) followed a sigmoidal relationship (Fig. 2b-c and Extended Data Fig. 6). Such normalization is physiologically relevant, since water movement depends entirely on relative differences in water potential rather than absolute values. A zero-one inflated beta regression model explained 91% of the variance in the data set, indicating high predictive power of local water potentials for lateral root patterning.

Our regression model allowed us to make predictions regarding how biophysical properties of the root and its environment might impact hydropatterning. A set of predicted lateral root distributions under various parameter perturbations is summarized in Extended Data Fig. 7. To facilitate the use of the model by non-specialists, we have generated an R Shiny app that allows the full parameter space to be explored through an interactive GUI (https://nrobbins.shinyapps.io/20161105_hydropatterning_app/).

Given the strong correlation between growth and competence for hydropatterning, we examined the predicted effects of altering growth dynamics on branching pattern using our model (Fig. 2d). Interestingly, the frequency of lateral root initiation toward air increased as the end of the growth zone was shifted towards the root tip and away from the competent/fixed-zone boundary. This was more pronounced at lower values of peak elemental growth rate, suggesting a synergistic effect between the two factors. We note that these simulations were performed assuming constant position and size of the competent zone, which allowed for uncoupling of growth from competence. Contrastingly, the effect of growth zone positioning on lateral root patterning was strongly reduced when the competent zone was configured to track with the growth zone (Fig. 2e). This indicated that tight coordination between growth as a signal generator and competence as a signal receiver were likely to be important for hydropatterning.

Based on these simulations, we hypothesized that the ability of the root to locally distinguish regions of high and low water availability may depend on the overall rate of growth-sustained water uptake in the competent zone. To explore this question further, we scored lateral root patterning in seedlings exposed to different growth inhibitors. Seedlings were treated with sodium orthovanadate (Na_3_VO_4_) and diethylstilbestrol (DES), two inhibitors of plasma membrane H+-ATPases which partly function to acidify the cell wall and promote wall-loosening expansin activity during cell elongation^8,18–20^. Consistent with our hypothesis, hydropatterning was disrupted in treatment conditions that also reduced growth (Fig. 3a-b). Comparable results were obtained using citric acid, which increases pH-buffering capacity of the external medium, as well as low-temperature stress (Fig. 3c-d). The predictions of our regression model significantly correlated with empirical observations of lateral root patterning in this data set, with correlation coefficients between 0.62 and 0.95 depending on treatment condition (p < 0.0003; Fig. 3e). This variation is suggestive of treatment-specific effects on lateral root patterning that are independent of alterations to growth dynamics. Nonetheless, these observations provide validation of the predictive power of the model under a broad range of conditions, and support our hypothesis that growth is required for lateral root hydropatterning.

**Figure 3.**
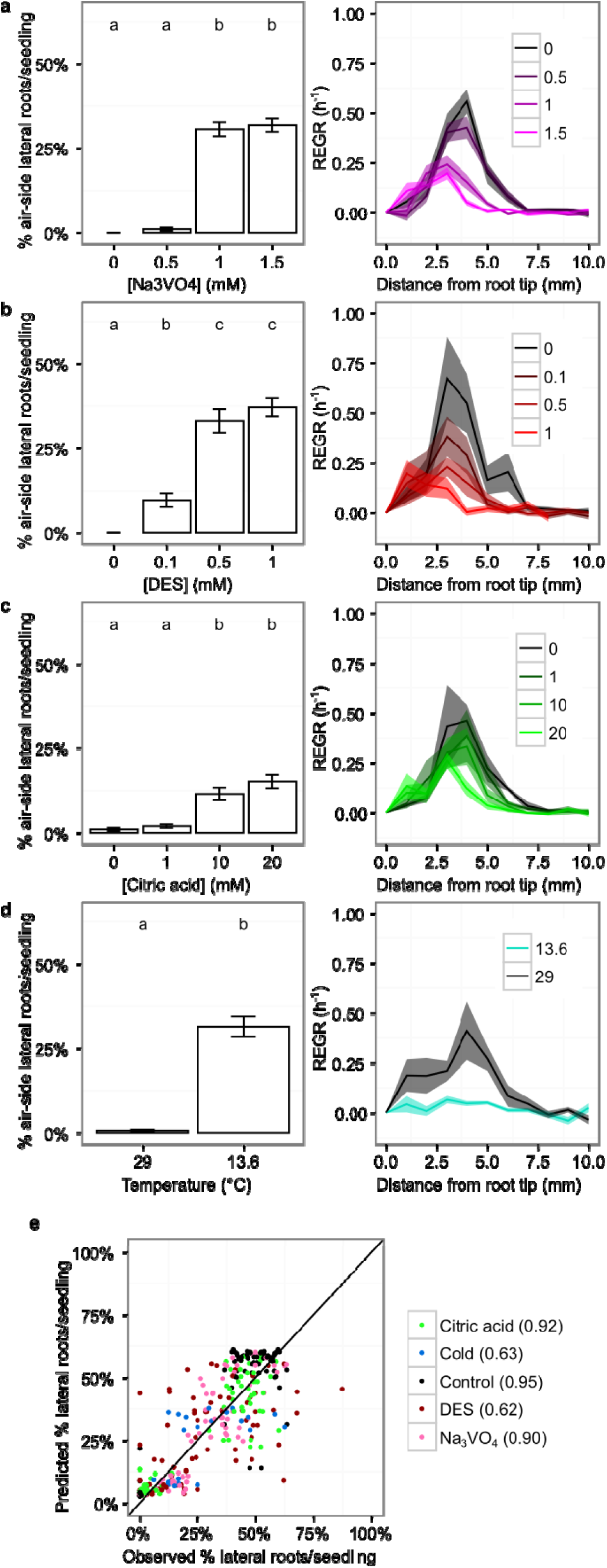
Growth is necessary for lateral root hydropatterning. **a-d**, Average frequency of air-side lateral root initiation (left) and relative elemental growth rate (REGR) (right) in indicated treatment condition. Primary roots were grown between two agar sheets, with chemical treatments applied to both agar-contacting sides. Lateral root emergence was quantified in tissues exposed to air gap between agar sheets. Error bars and shaded regions, standard error of the mean. Significantly different groups denoted with different letters (p < 0.05). N = 10-16 and 7-8 seedlings for lateral root distribution and REGR, respectively, across 2 experiments per treatment level. **e**, Observed and model-predicted lateral root distributions for samples used for kinematic growth analysis in **a-d**. Diagonal line denotes perfect prediction of empirical data by the model. Values in parentheses denote Pearson’s correlation coefficient for comparison of empirical and predicted values. Each coefficient was significantly different from 0 (p < 0.0003).

We asked whether growth-sustained water potentials had a broader impact on cellular signaling and physiology by performing transcriptomic profiling of longitudinal domains of the root corresponding to the competent and fixed regions, and separately profiling tissue in contact with air or agar (Fig. 4a, Supplementary Table 1, and Gene Expression Omnibus database accession no. GSE92406). We detected a total of 25,835 unique transcripts, 1,559 of which were significantly differentially expressed between the air- and agar-exposed sides of the root. Of these, 1,461 were differentially expressed in the fixed zone, suggesting that the functional divergence of the two sides occurred primarily after competence was lost.

**Figure 4.**
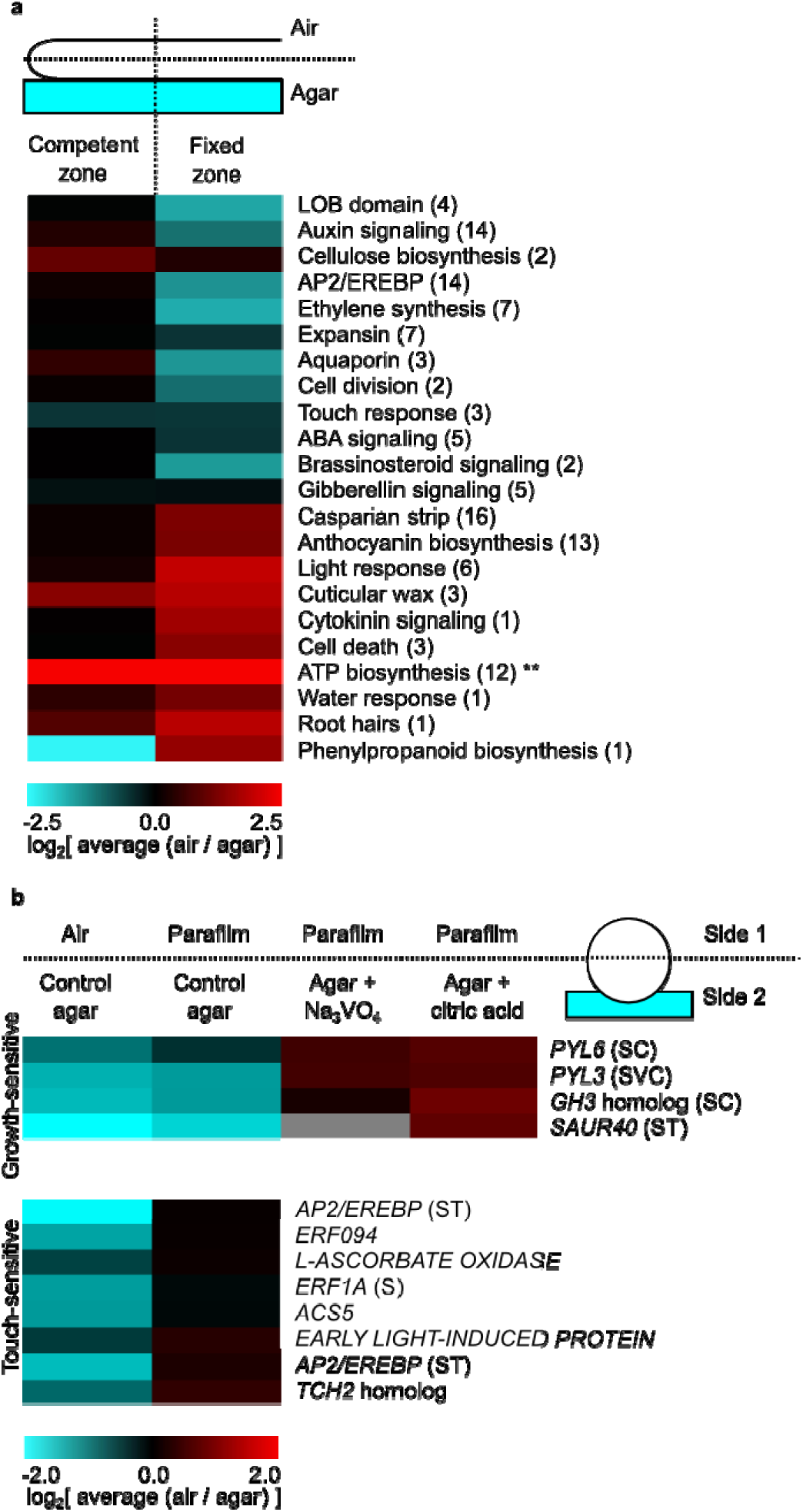
Growth plays a role in regulation of gene expression by water availability. **a**, Expression patterns of side-biased gene categories identified by RNA-seq. Seedlings were grown along a single agar surface and sectioned according to diagram (top). Air/agar FPKM ratio was computed for each gene, and averaged by category. Resulting values were then log-transformed. Number of genes per category indicated in parentheses. **, values for ATP biosynthesis were outliers (6.34 and 6.77 for competent and fixed zones, respectively), and were represented as 2.5 for plotting purposes. **b**, Expression patterns of a subset of genes identified in a measured by RT-qPCR. Seedlings were grown between agar and indicated low-water availability substrate (top). Agar was supplemented with 1.5 mM Na_3_VO_4_ or 20 mM citric acid where indicated. Air/agar relative expression ratio (arbitrary units) was computed for each gene and log-transformed. Parentheses denote genes identified as statistically significantly regulated (p < 0.05): S, side-biased in air/control agar and Parafilm/control agar; T, touch-sensitive; V, Na_3_VO_4_-sensitive; C, citric acid-sensitive. Growth-sensitive and touch-sensitive gene clusters identified based on hierarchical clustering. Data shown for fixed-zone tissues.

The side-biased transcriptome included a number of pathways known to be affected by hydropatterning, including anthocyanin biosynthesis, root hair development, programmed cell death, lignin accumulation, and signaling associated with the plant hormone auxin^1,21^. Several agar-induced genes encoded lateral organ boundaries (LOB) domain transcription factors, including ROOTLESS CONCERNING CROWN AND SEMINAL LATERAL ROOTS (RTCS)^22^. RTCS homologs in Arabidopsis and rice are involved in early stages of lateral root development^23–26^, suggesting that these genes may serve as markers of the process in maize. Our analysis also revealed novel pathways regulated during hydropatterning, including brassinosteroid and ethylene signaling, and water transport.

In order to test the role that growth has in determining these gene expression patterns, we used high-throughput RT-qPCR to quantify expression of a panel of side-biased genes in seedlings treated with Na_3_VO_4_ or citric acid (Fig. 4b and Supplementary Table 3). To determine if differentially expressed genes were responsive to water availability or mechanical contact, we compared roots exposed to air and agar to roots grown between agar and a sheet of Parafilm. A subset of the genes examined were equally induced by Parafilm and control media, indicating that mechanical contact alone was likely responsible for their induction by agar (touch-induced). Amongst the genes that were induced by agar and not touch alone, we identified several that lost their water-biased expression under growth inhibition. Within this set were genes in the auxin pathway, as well as members of the PYRABACTIN RESISTANCE1/PYR1-LIKE (PYL)/REGULATORY COMPONENT OF ABA RECEPTORS family of receptors for abscisic acid, a hormone broadly involved in responses to water-deficit stress^27^. This observation demonstrates that a portion of the water-responsive transcriptome is sensitive to changes in growth dynamics, providing evidence of a more general role of growth in the response of the root to water availability.

Together, our data support a model in which growth-associated water uptake, in conjunction with spatial heterogeneity in local water availability, generates internal gradients of tissue water potential that inform developmental patterning and gene expression. Thus, growth creates a physical state in which water perception can occur. This “sense-by-growth” mechanism illustrates that the perception of water is dependent on a state of disequilibrium established by the organism that allows meaningful spatial information to be derived from the external environment. Our work thus represents a significant advance in our understanding of the processes governing water sensing in plants, knowledge of which will be integral to improving crop water-use efficiency to meet the demands of a growing world population.

## Methods

### Plant materials and growth conditions

Experiments were performed with wild-type Zea mays inbred B73 provided by E. Vollbrecht. Seeds were sterilized using a heat/bleach treatment and germinated on autoclave-sterilized 100-lb germination paper (Anchor Paper Company) soaked with autoclave-sterilized deionized water for 2 days^1,28^. Germinated seedlings were transferred to autoclave-sterilized agar media containing 1/2X Murashige & Skoog basal salts (Caisson Labs) and 0.5 g/l MES hydrate (Sigma-Aldrich) in 120 * 120 * 17-mm square plastic plates (USA Scientific). Positions of primary root tips at the time of agar transfer were marked on the backs of the plates with a marker. 1% (w/v) agar (BD, Difco) was used for radial sectioning and RNA-seq experiments, and 2% agar was used for all others. Where indicated, seedlings were grown between two agar sheets as done previously^1,28^, with the exception that no blot paper was placed between the roots and media. For RT-qPCR experiments, a square of Parafilm M^®^ (Bemis Company, Inc.) was placed between the root and one agar sheet to induce a touch response without supplying available water. Where indicated, media was supplemented with citric acid, monohydrate (Sigma-Aldrich); diethylstilbestrol, mixture of cis and trans (Sigma-Aldrich); mannitol (PhytoTechnology Laboratories); or sodium orthovanadate (Sigma-Aldrich). Diethylstilbestrol was added from a 200-mM stock in DMSO. Sodium orthovanadate was added from a 200-mM stock in water, prepared by repeatedly heating and decreasing the pH of the solution with hydrochloric acid until it stabilized as a clear solution at pH 10.0. Citric acid, mannitol, and sodium orthovanadate were added prior to media autoclaving, while diethylstilbestrol was added immediately afterwards. To control for the presence of DMSO in experiments conducted with diethylstilbestrol, equal volumes of DMSO were added to all media conditions. Agar plates were incubated vertically in a growth chamber (Percival, CU41L4) under 16 h light (29°C, 100μE light) / 8 h dark (24°C) cycle. For low-temperature experiments, chamber temperature was set to a constant 13.6°C, and light levels in both the low- and high-temperature conditions were set to 20 μE.

### Kinematic growth analysis

Methods for kinematic analysis were adapted from prior studies^4–6^. 1 day following transfer to agar media, India ink (Higgins) was dotted along the primary root between 0 and ~10 mm from the tip. Where necessary, agar sheets were temporarily removed from seedlings to allow ink to be applied. Roots were then photographed with a digital camera (Kodak EasyShare C533 or Nikon D5000) every 20 min for ~3 h. Image analysis was performed in Fiji^29^. Images were aligned in Fiji using the Linear Stack Alignment with SIFT plugin^30^. X-Y coordinates of the root tip and the proximal and distal edges of each ink mark were recorded using the Manual Tracking plugin. Data analysis was done in R using the plyr library^31,32^. Displacement velocity for each tracked mark was calculated as the slope of a linear regression of position against time fitted to all data points for that mark, with position defined as the distance from the root tip. These velocity values were paired with mark positions at time 0 to generate a plot of velocity against distance from the root tip. These data were binned by averaging values within bins of 0.5 mm. Relative elemental growth rate was calculated by numerically differentiating the velocity-distance curve with respect to distance. For samples used in modeling tissue water potential, positions with missing data were inferred as having the same relative elemental growth rate as the next known value in the curve. The end of the growth zone was defined as the first point after the peak elemental growth rate where the rate fell below 0.01 h^-1^.

### Competent zone determination

To determine the competent/fixed zone boundary, seedlings were grown along a single agar surface for 1 day, then divided into 4 treatment conditions: 1, agar sheets were applied to the air-exposed side of the primary root; 2, ink was applied to the primary root and kinematic growth analysis was performed; 3, both ink and agar sheets were applied and kinematic analysis was performed; and 4, no perturbations. Seedlings from all treatments were photographed immediately after ink/agar application with a dissection microscope (Leica M165 FC microscope with KL 1500 LED light source and DFC 500 camera), and again once lateral roots emerged 3-4 days afterwards. Marks on the backs of the plates were used as fiducial markers to align images from the two time points. For seedlings with applied agar sheets, a line was drawn between branching and non-branching regions of the primary root in the final image, and traced horizontally to the initial image (Extended Data Fig. 1a). The distance from this line to the root tip was taken as the position of the competent/fixed boundary. Statistical comparisons of the positions of the competent/fixed boundary and the end of the growth zone were done using a mixed-effects analysis of variance (ANOVA). Landmark (competent/fixed boundary or end of growth zone), experiment ID, and their interaction were used as predictors of position measurements, with plant ID included as a random effect. Post-hoc analysis was performed using a general linear hypothesis test to determine the average and standard error of the difference in position between the two landmarks.

To determine the size of the competent zone, seedlings were grown as above and divided into 3 treatment conditions: 1, agar sheets were applied to the entire length of the air side, with the bottom edge flush with the root tip; 2, agar sheets were applied to the air side and manually cut to leave a thin strip ≤5 mm in length at the root tip; 3, no agar sheets applied. Seedlings were photographed using a dissection microscope 3 days after agar application. Applied agar sizes were measured using images in Fiji^29^. Seedlings were categorized as 0% or 100% to denote 0 or ≥1 lateral roots emerged on the treated side, respectively. The size of the competent zone was determined as the minimum-sized sheet of agar that induced lateral roots. A generalized linear regression (binomial family, logit link) was performed to predict probability of lateral root induction based on sheet length, experiment ID, and their interaction. Data analysis was done in R using nlme, multcomp, and tidyr libraries^31,32–35^.

### Water potential modeling

The growing maize primary root was modeled as a right circular cylinder divided into longitudinal segments. Each segment was composed of 3 concentric tissue layers: epidermis, cortex, and stele, with diameters of 0.1, 0.5, and 0.4 mm, respectively (Extended Data Fig. 2a). Dimensions were determined from radial sections of a maize primary root, inbred B73 (e.g. Fig. 1b). The epidermis and cortex were further divided into quarters, yielding 9 unique compartments per segment.

The height of each segment was assumed to increase at a rate defined by the local elemental growth rate (changes in root width are not taken into account). 90% of the resulting change in volume over time was assumed to be due to water uptake. Thus, the rate of water uptake for each segment was calculated as: J_V_ = 90% * π * r^2^ * h * REGR, where J_V_ is volumetric water flow rate (m^3^ s^-1^), r is root radius (0.5 * 10^-3^ m), h is the initial height of the segment (distance between adjacent points in relative elemental growth rate plot, typically 0.5 * 10^-3^ m), and REGR is the relative elemental growth rate of the segment (h^-1^) (Extended Data Fig. 2b).

Sources of water were included on all exterior faces of the root, as well as an internal phloem source at the center of the root. Phloem was modeled as a cylinder 0.3 mm in diameter within the stele layer. The water potential and hydraulic conductivity of each source were independently specified. Source potentials were assumed to be −0.1 MPa, except in the case of mannitol-supplemented agar, where it was calculated as: −0.1 MPa - (400 / c), with c denoting mannitol concentration (mM). Hydraulic conductivities (m^3^ m^-2^ s^-1^ MPa^-1^) were 10^-5^ for 1% agar media, 2.5 * 10^-6^ for 2% agar media, and 3.66 * 10^-9^ for air^36,37^. Phloem conductivity was set to yield ~25% of total root water uptake as phloem-derived in all simulations^38^.

In order to calculate compartment water potentials, the network of compartments was modeled as an electric circuit (Extended Data Fig. 2c). Water was allowed to flow between adjacent compartments according to the equation: J_V_ = ΔΨ_w_ * Lp * A, where ΔΨ_w_ is the water potential difference between the compartments (MPa), Lp is hydraulic conductivity of the tissue (1.15 * 10^-7^ m^3^ m^-2^ s^-1^ MPa^-1 16^), and A is the surface area of the interface between the compartments (m^2^). Conductivities for movement from water sources to root compartments were calculated by treating root and source conductivities as being in series: Lp_source to root_ = (Lp_source_ * LP_root_) / (LP_source_ + LP_root_). Net water uptake rates for each compartment were calculated as a percentage of the uptake for the entire root segment based on the relative sizes of the compartments. Given these values and the potentials of the water sources, the rates of water movement between each compartment were calculated using a system of linear equations derived using Kirchhoff’s circuit laws (Extended Data Fig. 2d). These values were then used to calculate the water potentials of each compartment using a second system of linear equations derived from the relationship defined above (J_v_ = ΔΨ_w_ * Lp * A). All calculations for the model were performed in R and used the plyr library^31,32^.

The above method was adapted to Glycine max hypocotyls (Extended Data Fig. 3) with the following modifications. Each hypocotyl segment was divided into 9 concentric tissue layers with diameters that matched the positions where water potential was empirically measured ^13^. External water sources were excluded from the model. Water potential and conductivity of the internal water source were set to −0.06 MPa and 1 m^3^ m^-2^ s^-1^ MPa^-1^, respectively. Several values of tissue hydraulic conductivity were examined. For each conductivity value, the water potential estimates at the point of maximal elemental growth rate were compared to empirical values. A t-statistic was calculated at each point in the water potential profile: t = (empirical mean - estimated value) / (empirical 95% confidence interval / 1.96). Two-tailed p-values were then determined for each t-statistic (degrees of freedom = 11, based on median sample size for empirical water potential measurements). Cumulative p-values for each conductivity condition were calculated using Fisher’s combined probability test.

Example R scripts and input files used to calculate tissue water potential are available through a Github repository (https://github.com/nerobbin/20161214hydropatterning).

### Lateral root quantification

Lateral roots from seedlings 6-7 days after seed imbibition were counted in four quadrants of the primary root (two contacting agar and two exposed to air). Counts in the two air-exposed quadrants were summed together, giving three total categories. Counts were done from the position of the primary root tip at the time of transfer to agar media to the bottom edge of the applied agar sheet. For each seedling, counts in each category were converted to percentages by dividing by the total count for that seedling. For data from mannitol-treated seedlings (Extended Data Fig. 4c), statistical analysis was performed using a mixed-effects ANOVA. The model used treatment condition, lateral root category, experiment ID, and all two- and three-way interaction terms as predictors for lateral root percentages, with seedling ID as a random effect. A condition:category interaction term with p < 0.05 indicated a significant effect of the treatment on lateral root patterning. Post-hoc comparisons were performed using a general linear hypothesis test. Matching categories were compared pairwise between the different treatment conditions. Treatments that significantly differed in any one of the three categories (p < 0.05) were deemed to have significantly different lateral root distributions, and were assigned to different significance groups. For data from growth-inhibitor-treated seedlings (Fig. 3a-d), statistical analysis was performed using a two-way ANOVA. The model used treatment condition, experiment ID, and their interaction term as predictors for air-side lateral root percentages. A condition term with p < 0.05 indicated a significant effect of the treatment on lateral root patterning. Post-hoc analysis was performed using Tukey’s honest significant difference test, with significance groups assigned using a significance threshold of p < 0.05. Analysis was performed in R using nlme, multcomp, and tidyr libraries^31,33–35^.

### Regression analysis

The regression model relating empirical lateral root distributions and modeled tissue water potentials was trained using data from mannitol-treated seedlings for which both growth-kinematics and lateral-root data were available (Extended Data Fig. 4). Growth curves from individual seedlings were used to estimate tissue water potentials. For each root quadrant, the sum of the potentials in the epidermis layer in the competent zone (4.0-5.5 mm from the tip, Fig. 1e and Extended Data Fig. 5b) were plotted against respective lateral root percentages. Epidermal potentials were used based on evidence implicating outer tissues as important sites of early hormonal signaling events upstream of water-induced lateral root initiation^1^. Water potentials were normalized in two ways: by dividing each quadrant potential by the sum of all quadrant potentials for that seedling (% water potential), or by mean-centering the potentials from each seedling. Data from both normalization methods were used to fit zero-one inflated beta models in R using the gamlss library^31,39^. This type of model was chosen because the dependent variable was proportion data with many values at 0% or 100%. Variance explained was assessed using a pseudo-R^2^ value: 1 - [∑((observed value - predicted value)^2^) / Σ ((observed value - mean of observed values)^2^)]. The optimal model was selected based on pseudo-R^2^, root-mean-square error, Akaike Information Criterion, and residuals plots. Model validation was done by comparing model-predicted values to independent empirical data generated from seedlings under growth inhibition. For each treatment, Pearson’s correlation coefficient was calculated comparing empirical and predicted data, and statistical significance of the correlation was tested using the cor.test function in R^31,39^.

Data sets for training and validation of the regression model are available through a Github repository (https://github.com/nerobbin/20161214hydropatterning).

### Predictive model and Shiny app

A predictive model relating growth and biophysical properties of the root and its environment to lateral root patterning was written in R as a Shiny app^31,40^. A simulated growth curve is used to generate water potential estimates. The curve is a parabola fitted to user-defined values for the start and end of the growth zone, and the peak elemental growth rate positioned at the center of the growth zone. Growth rates are determined at 0.5-mm intervals along the curve. The curve is displayed using the ggplot2^41^ and scales^42^ libraries. Water potentials are displayed as a choropleth map using the Cairo^43^, ggplot2^41^, ggmap^44^, maptools^45^, and rgeos^46^ libraries. The regression model equation in Extended Data Fig. 6c is used to calculate predicted lateral root distributions, which are displayed as a bar graph. Each parameter value is freely modifiable, with the following exceptions: external water source conductivities (2.5 * 10^-6^ and 3.66 * 10^-9^ m^3^ m^-2^ s^-1^ MPa^-1^ for agar and air, respectively); relative diameters of root tissue layers (scaled proportional to values for maize primary root); and parameters governing the regression model equation.

### RNA-seq

Primary roots of seedlings 3 days post-imbibition were manually dissected using an X-ACTO^®^ #1 precision knife with #11 blades under a dissection microscope (Olympus SZ61 with Schott ACE light source) as shown in Fig. 4a. Competent- and fixed-zone tissues were taken from 0-5 and 5-15 mm from the root tip, respectively. 3 biological replicates were collected for each tissue type, with each replicate composed of tissue pooled from 2 seedlings. Tissues were flash-frozen in liquid nitrogen and stored at −80°C until processed. RNA was isolated using the ZR Plant RNA MiniPrep™ kit (Zymo Research). Tissues were homogenized using a FastPrep^®^-24 (MP Biomedicals), twice for 30 s at 6.0 m/s. On-column DNase digest was done using RNase-Free DNase Set (50) kit (QIAGEN). RNA yields were quantified using the Qubit^®^ RNA HS Assay on a Qubit^®^ 2.0 Fluorometer (Thermo Fisher Scientific). Ribosomal RNA was removed using the RiboZero™ Magnetic Kit for Plant Seed/Root (Epicentre), followed by cleanup using the Zymo RNA Clean & Concentrator™-5 kit (Zymo Research). Sequencing libraries were prepared using the NEBNext^®^ Ultra™ Directional RNA Library Prep Kit for Illumina^®^ (New England Biolabs). Library concentrations were measured using the Qubit^®^ dsDNA HS Assay (Thermo Fisher Scientific), and quality was assessed using an Agilent 2100 Bioanalyzer (Agilent Technologies). Libraries were pooled based on concentrations assessed by qPCR on a LightCycler^®^ 480 (Roche Life Science) using primers directed against adapter sequences. The library pool was sequenced on a HiSeq 2000 (Illumina) (paired reads, 2x101-bp plus index) through Stanford Center for Genomics and Personalized Medicine. One library (contact-side competent-zone replicate 3) was run on a separate flow cell due to an error during library preparation.

Post-filter read files were uploaded to the iPlant Discovery Environment for preprocessing: Quality was assessed using FastQC v0.10.1^47^, Scythe v0.981^48^ and Cutadapt v1.3^49^ were used to remove detected adapter sequences, and Sickle v1.0^50^ was used for quality trimming (quality format = sanger). Files with paired sequences were uploaded to the Galaxy Project website to perform the following: Sequences were mapped to the Zea mays B73 genome (AGPv3, Ensembl 21^51^) using TopHat v2.0.9^52^, and transcripts were assembled using Cufflinks v2.1.1 and Cuffmerge^53,54^. BAM files were converted to sorted SAM files using Samtools v0.1.19^55^, and transcript counts were calculated using HTSeq 0.5.4p3^56^. Differential expression was assessed using Cuffdiff v2.2.1^57^ and DESeq2 v1.14.0^58^. Pairwise comparisons were made between air- and agar-side tissues. Competent and fixed tissues were analyzed separately. For DESeq2, separate analyses were performed with grouping between samples from matching biological replicates taken into account or not taken into account. Significance was determined with false discovery rate-adjusted p-value < 0.05 for all comparisons. Functional annotation and Arabidopsis orthologs for significantly differentially expressed genes were retrieved using gProfiler^59^. For the top 25 up-regulated genes in each tissue section, missing annotation data were supplemented by running a BLAST search on mRNA sequences downloaded from MaizeGDB^60^. Genes were manually assigned to functional categories based on annotation information. Fold-changes of all categorized genes were calculated as air-side FPKM / contact-side FPKM, and averaged across the biological replicates. These values were then averaged by category, log2-transformed, and plotted as a heat map in Multiple Experiment Viewer v4.8.1^61^. Hierarchical clustering was done using Pearson correlation metric.

### RT-qPCR

Seedlings were grown in conditions shown in Fig. 4b. Root dissection and RNA isolation were done as for RNA-seq. 3 biological replicates were prepared per condition. Genomic DNA contamination was assessed by performing endpoint PCR on purified RNA using IDP7742 primers (Supplementary Table 3). cDNA was synthesized using iScript™ Reverse Transcription Supermix for RT-qPCR (Bio-Rad). Gene expression was profiled using a Biomark™ HD 48.48 Dynamic Array™ (Fluidigm) according to manufacturer specifications. Primers were synthesized by IDT (Supplementary Table 3) and tested by endpoint PCR on seedling leaf genomic DNA using GoTaq^®^ Green Master Mix (Promega). Genomic DNA was purified using ZR Plant/Seed DNA MiniPrep™ (Zymo Research). A subset of primer-pairs yielded multiple PCR products in this reaction and were removed from downstream analyses.

Data analysis was performed in R with tidyr library^31,35^. Transcript abundances were calculated based on normalized fluorescence intensity plots^62^. Reactions for which no Ct value was called by the Biomark™ software were omitted. Within each sample, abundances were normalized by dividing by the abundance of the loading control assay. The control (GRMZM2G015295_T03, encoding peptide chain release factor subunit 1) was selected based on its high and uniform expression amongst all samples in the RNA-seq experiment. Normalized abundances were used for statistical analysis. Each gene and zone of the root were analyzed separately, and only biological replicates that had expression data from both sides of the root were used. First, a mixed-effects ANOVA was performed on data pooled from air/control-agar and Parafilm/control-agar conditions. The model tested for effects of side, condition, and their interaction, with replicate ID as a random effect. Genes with a significant side term (p < 0.05) were deemed side-biased, and those with a significant interaction term were deemed touch-sensitive. For genes that were side-biased but not touch-sensitive, the ANOVA was repeated with data from either the Parafilm/Na_3_VO_4_-agar or Parafilm/citric acid-agar conditions included. If the interaction term was significant in either assay, then the gene was deemed growth-sensitive. Average fold-changes were plotted and clustered as described for RNA-seq data.

## Acknowledgements

We thank Robert E. Sharp, John S. Boyer, Kenneth A. Shackel, and Wendy K. Silk for inspiration and suggestions on experimental design. We also thank Silk and members of the Dinneny lab for critical evaluation of the content of the manuscript. Funding was provided by the Carnegie Institution for Science Endowment to JRD. Research reported in this publication was supported by NIGMS of the National Institutes of Health under award number T32GM007276. The content is solely the responsibility of the authors and does not necessarily represent the official views of the National Institutes of Health. This material is based upon work supported by the National Science Foundation Graduate Research Fellowship under Grant No. DGE-1147470. Any opinion, findings, and conclusions or recommendations expressed in this material are those of the authors(s) and do not necessarily reflect the views of the National Science Foundation.

## Author Contributions

N.E.R. and J.R.D. designed the study and wrote the manuscript. N.E.R. carried out experiments and performed data analysis.

## Author Information

RNA-seq data were deposited to the Gene Expression Omnibus database, accession no. GSE92406. Raw data and R scripts for water potential estimation and regression analysis were uploaded to a Github repository (https://github.com/nerobbin/20161214hydropatterning). The authors declare no competing financial interests. Correspondence should be addressed to J.R.D.

---

**Extended Data Figure 1.**
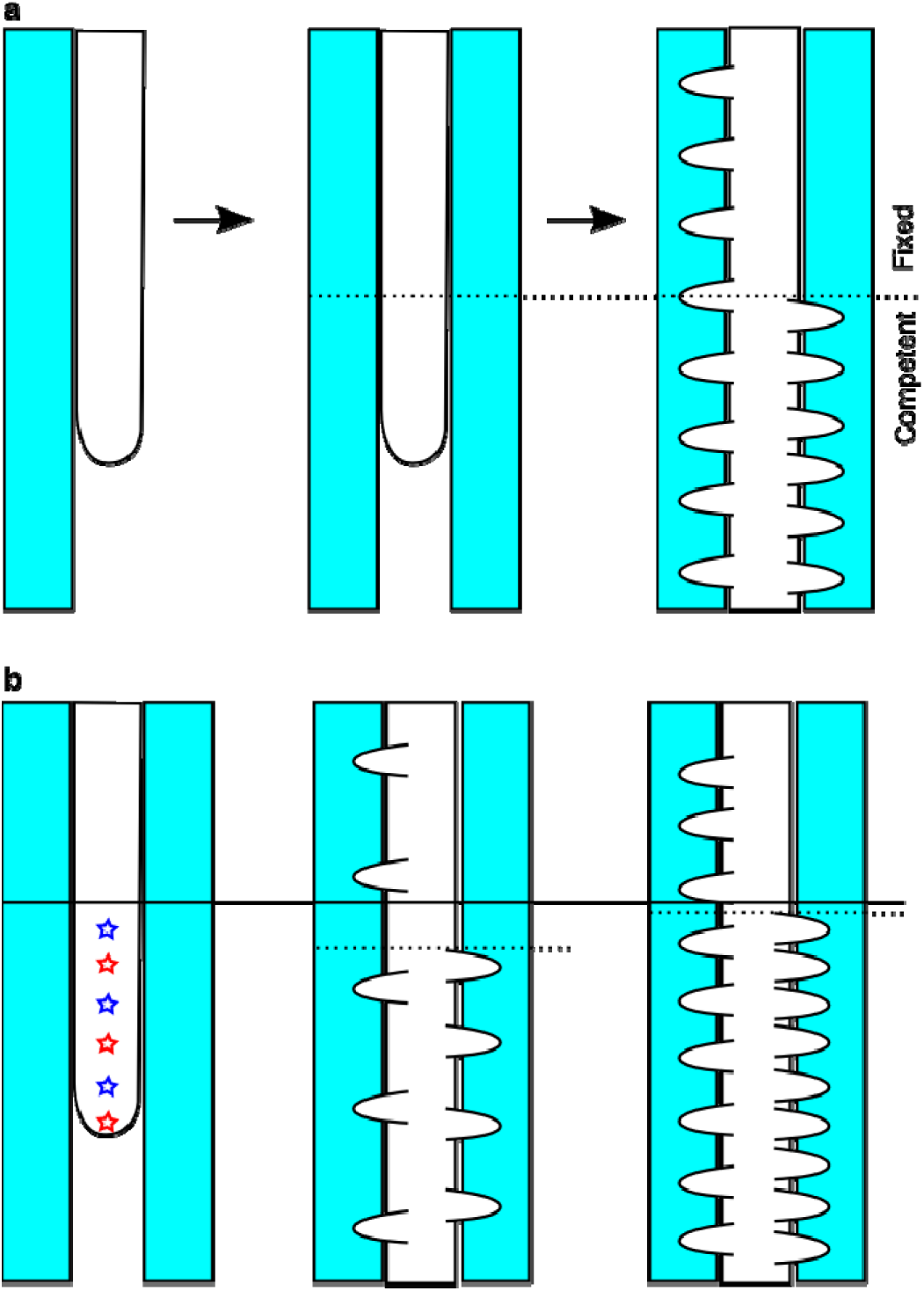
Method to determine the location of the competent/fixed boundary for lateral root hydropatterning. **a**, A primary root is grown along an agar surface (left), an agar sheet is applied to the air side (center), and lateral roots emerge several days later (right). Competent and fixed regions are determined based on the presence or absence of lateral roots toward the applied agar, respectively. A horizontal line drawn from the right to the center image determines the initial position of the competent/fixed boundary. **b**, Lateral root density affects spatial resolution of this method. The true position of the competent/fixed boundary is shown (solid line), and stars denote possible positions of lateral root emergence towards the applied agar (left). If lateral roots are sparse (emergence at red stars), a discrepancy between the true and observed boundary is introduced (dotted line, center). This is minimized with increasing density (emergence at all stars, right).

**Extended Data Figure 2.**
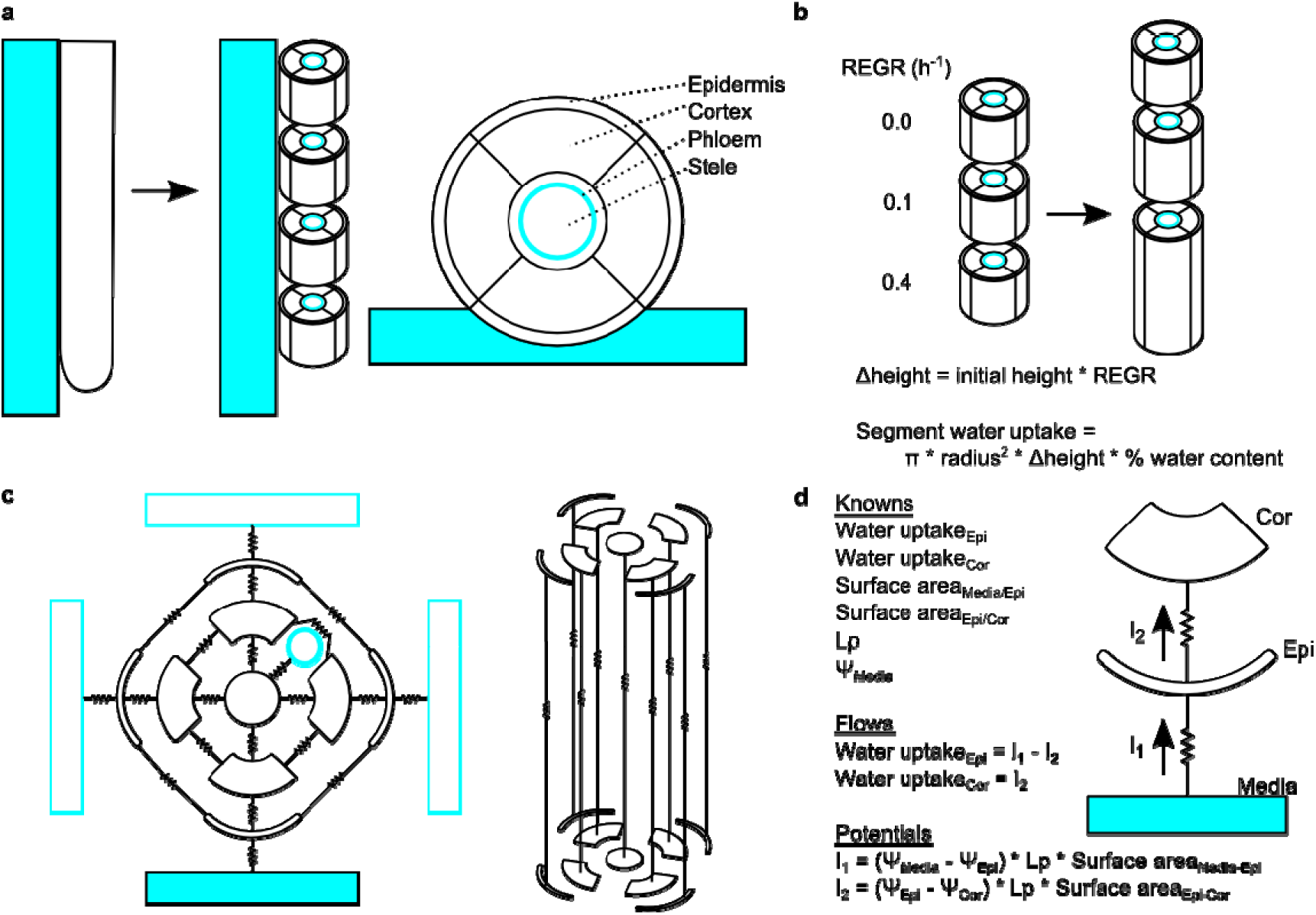
Pictorial explanation of mathematical model used to estimate tissue water potentials resulting from growth. **a**, The growing organ is treated as a series of right circular cylinders, each divided intocompartments. Water is allowed to flow between adjacent compartments both radially and longitudinally, and can be taken up from the external environment or provided internally via the phloem. **b**, Growth is modeled as an increase in cylinder height over time, which is determined based on local relative elemental growth rate (REGR). A user-specified percentage of the change in cylinder volume is assumed to be due to uptake of water. **c**, The network of root compartments is treated as an electric circuit. Connections for radial water flow in one segment (left) and longitudinal flow between two segments (right) are shown. Cyan, water sources: filled box, agar; hollow box, air; circle, phloem. **d**, Example of calculations used to derive compartment water potentials. Compartment water uptake rates and surface areas of compartment interfaces are calculated based on compartment geometry and total segment water uptake (b). Hydraulic conductivity (Lp) and media water potential (Ψ_media_) are user-specified. Inter-compartment water flow rates (I, arrows) are first determined using a system of equations (“Flows”), and are then used to calculate compartment water potentials in a second system of equations (“Potentials”). Note that this example does not take into account media hydraulic conductivity, which modulates Lp for I_1_. See Methods for complete explanation.

**Extended Data Figure 3.**
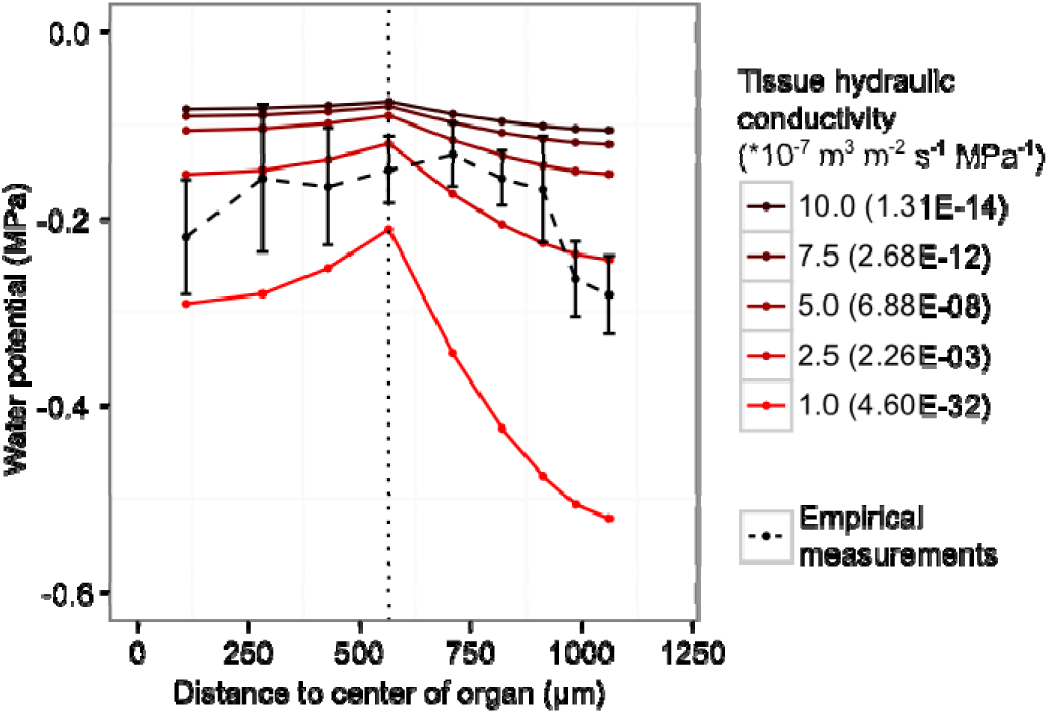
Comparison of mathematically modeled and empirically measured growth-sustained tissue water potentials along the radius of soybean hypocotyl. Empirical water potentials and tissue growth rates used to generate model estimates were derived from previously published data^13^. Estimated values were taken from the position of maximal relative elemental growth rate along the length of the hypocotyl. Vertical dotted line, position of water source in modeled organ. Error bars, 95% confidence interval. Values in parentheses indicate p-values for comparisons of estimated water potentials to empirical data at indicated hydraulic conductivity. Larger p-values indicate smaller overall deviations of estimates from empirical values. The maximal p-value is likely to occur at a conductivity between 2.5 and 5.0 * 10^-7^ m^3^ m^-2^ s^-1^ MPa^-1^.

**Extended Data Figure 4.**
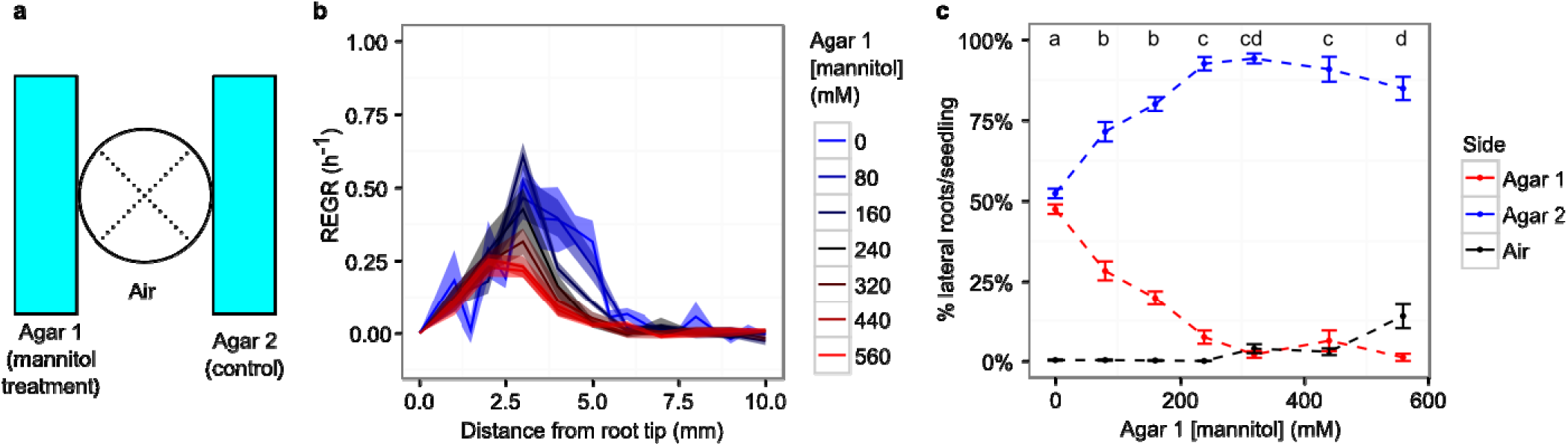
Growth kinematic analyses and lateral root patterning under mannitol treatment. **a**, Diagram of treatment conditions. Primary roots were grown between two agar sheets, with varying concentrations of mannitol applied in Agar 1. Agar 2 had 0 mM mannitol in all conditions. Dotted lines denote division of primary root for lateral root quantification. Counts on the two air-exposed sides were pooled together. **b-c**, Average relative elemental growth rate (REGR) profiles (**b**) and lateral root distributions (**c**) under indicated treatment conditions. Lateral root data are represented as averages of the percentage of lateral roots emerged on the indicated side of the primary root for each seedling. Error bars and shaded regions, standard error of the mean. Significantly different groups denoted with different letters (p < 0.05). Data are pooled from four experiments: two experiments examined 0, 80, 160, and 240-mM mannitol treatments, and the other two examined 0, 320, 440, and 560-mM mannitol treatments. Experiment-pairs were first analyzed separately to ensure no significant between-experiment variation existed. N = 8 seedlings for kinematic growth analysis and 15-16 for lateral root quantification for each condition in each experiment.

**Extended Data Figure 5.**
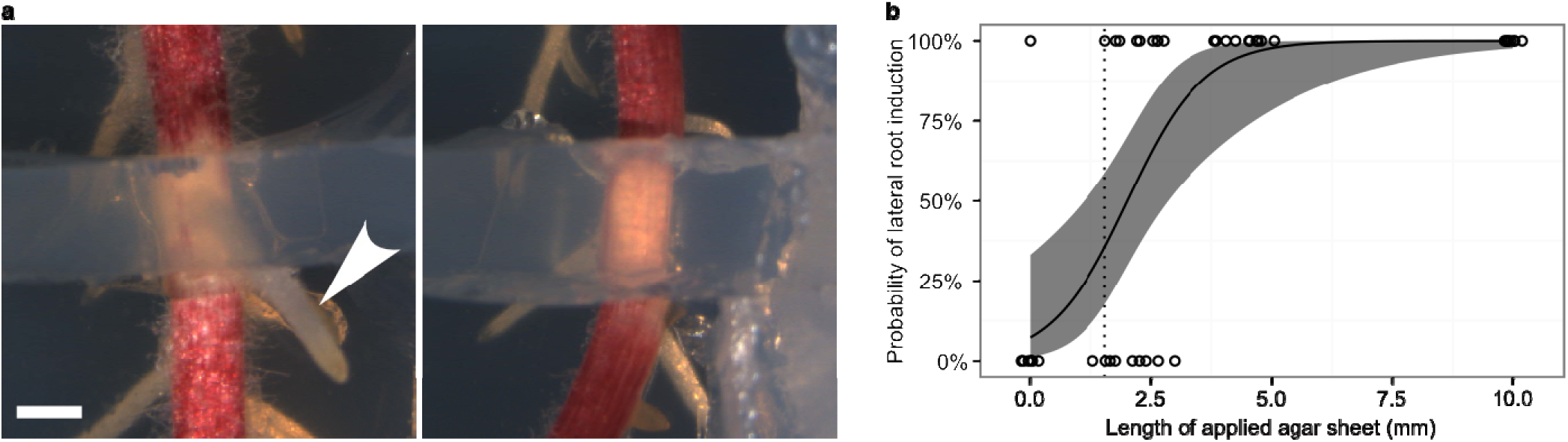
Determination of the root-apical boundary of the competent zone. **a**, Small agar sheets were applied to the air side of the primary root to determine the minimum sheet size necessary for lateral root induction. Representative images of sheets which have induced (left) or failed to induce (right) a lateral root (arrowhead) toward the applied agar are shown. Scale bar, 1 mm. **b**, Quantification of lateral root induction. Samples (circles) were categorized as 100% if applied agar induced lateral root development, or 0% if no induction was observed. Minimum inductive sheet size was 1.54 mm (dotted line). Logistic regression on these data provided estimates of the probability of induction with varying sheet size (curve). Sheet length was a significant predictor of lateral root induction in the regression model (p = 9.31 * 10^-6^). Shaded region, 95% confidence interval for regression curve. N = 47 across two experimental replicates. Data points at >10 mm were set to 10 mm for plotting purposes.

**Extended Data Figure 6.**
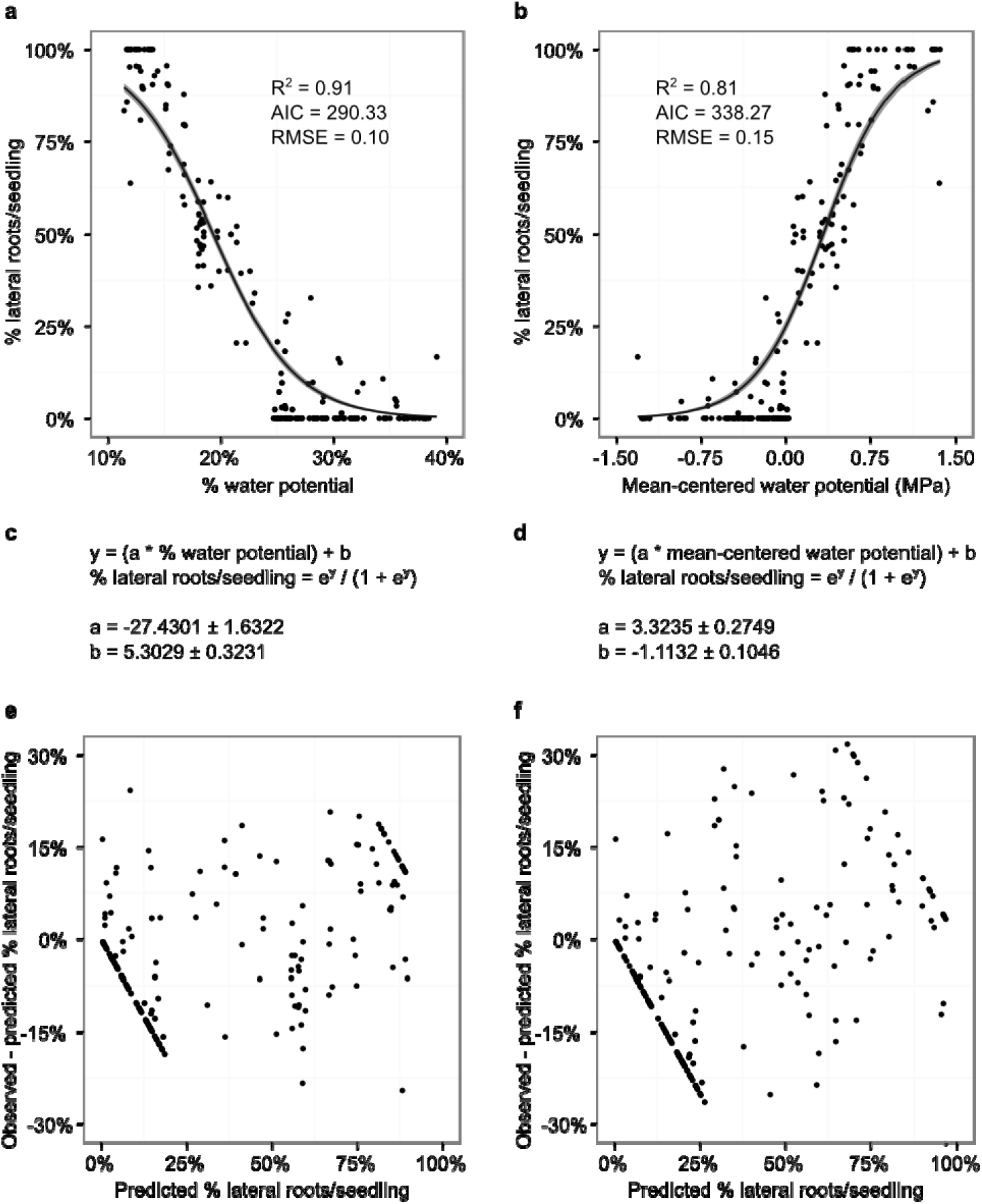
Comparison of two normalization methods for tissue water potential as predictors for lateral root patterning. **a-b**, Scatter plots of percent (**a**, reproduced from Fig. 2c) and mean-centered (**b**) water potentials against lateral root distributions for seedlings under mannitol treatment (Extended Data Fig. 4). Curves and shaded regions, means ± standard errors of best-fit lines for zero-one inflated beta regression models fitted to respective data sets. R^2^, pseudo-R^2^ value; AIC, Akaike information criterion; RMSE, root-mean-square error. **c-d**, Parameter estimates for models in **a-b**, respectively. Estimates of both parameter values significantly differed from 0 in both models (p < 2 * 10^-16^). **e-f**, Residuals plots for models in **a-b**, respectively.

**Extended Data Figure 7.**
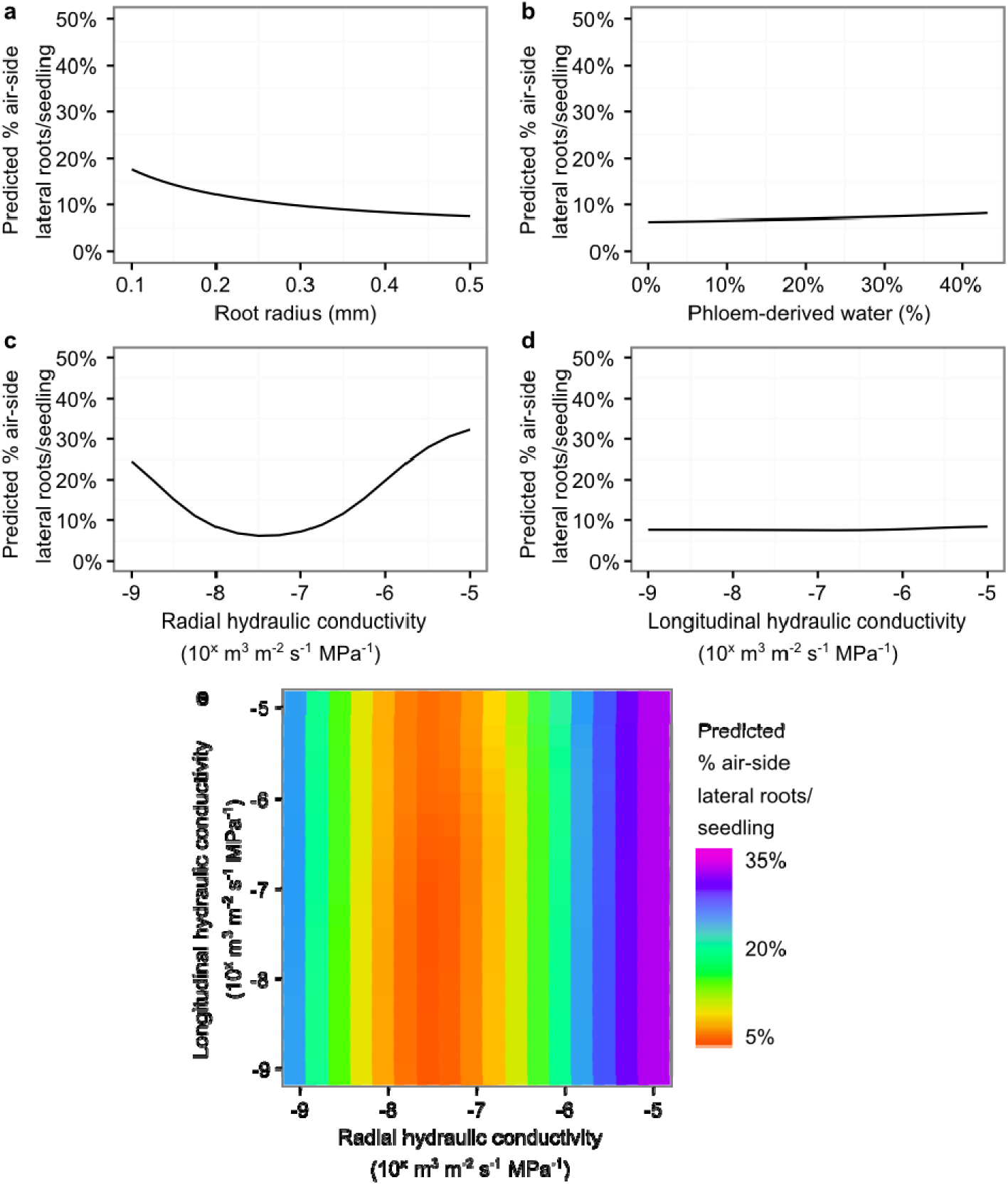
Modeling predicts that various biophysical parameters of the root influence its ability to pattern development in response to water availability. **a**, Decreases in primary root radius were associated with higher predicted rates of air-side lateral root initiation. This may explain the higher frequency of air-side lateral roots in Arabidopsis compared to maize, which has a narrower primary root radius^1^. **b**, Variation in % of total water uptake that is phloem-derived had relatively little effect on patterning. **c-d**, Changes in hydraulic conductivity in the radial direction (**c**), but not longitudinal (**d**), had an impact on lateral root patterning. This discrepancy may be due to the fact that longitudinal conductivity tends to alter absolute tissue water potentials without substantially altering relative water potentials. **e**, Model predictions with changes in both radial and longitudinal hydraulic conductivity. No synergistic effects between the two variables were observed. Unless otherwise indicated, parameter values were set to standards for maize primary root. All calculations were done using a simulated growth curve set to start at 0 and end at 5.5 mm from the root tip, with a peak relative elemental growth rate of 0.5 h^-1^.

